# Germination responses to changing rainfall timing reveal potential climate vulnerability in a clade of wildflowers

**DOI:** 10.1101/2023.03.22.533835

**Authors:** Samantha J. Worthy, Arquel Miller, Sarah R. Ashlock, Eda Ceviker, Julin N. Maloof, Sharon Y. Strauss, Johanna Schmitt, Jennifer R. Gremer

## Abstract

The timing of germination, driven by seasonal cues, is critical for the life cycle of plants. Variation among species in germination responses can reflect evolutionary processes and adaptation to local climate and can affect vulnerability to changing conditions. Indeed, climate change is altering the timing of precipitation and associated temperatures, which may interact with germination cueing to affect the timing, quantity, and speed of germination. Germination responses to change can then have consequences for individual fitness, population dynamics, and species distributions. Here, we assessed responses to the timing of germination-triggering rains and corresponding temperatures for 11 species spanning the *Streptanthus* (*s.l.*) clade (Brassicaceae). To do so, we experimentally manipulated the onset date of rainfall events, measured effects on germination fraction and rate, and evaluated whether responses were constrained by evolutionary relationships across the phylogeny. We then explored the possible consequences of these responses to contemporary shifts in precipitation timing. Later onset rains and cooler temperatures significantly reduced germination rates for all species. Germination fractions decreased with later rains and cooler temperatures for all but three *Caulanthus* species. Species’ germination responses to the timing of rainfall and seasonal temperatures were phylogenetically constrained, with *Caulanthus* species appearing less sensitive. Further, six species are likely already experiencing significant decreases in germination fractions or rates (or both) with observed climate change, which has shifted the timing of rainfall towards the cooler, winter months in California. Overall, our findings highlight the importance of the germination responses to seasonal timing, how they have evolved across the clade, and their implications under climate change.

## Introduction

The seasonal timing of seed germination depends critically on a species’ germination niche-the range of environmental conditions under which germination can occur. Germination timing in turn is critical to seedling establishment, seasonal phenology, and ultimately plant fitness (Kalisz, 1986; Donohue et al., 2010; Gremer et al., 2020). Precipitation and temperature are critical cues for germination (Probert, 2000; Finch-Savage and Leubner-Metzger, 2006; Burghardt et al., 2015; Barga et al., 2017; Puglia et al., 2018), in particular, the timing of precipitation interacts with temperature associated with rain events to drive germination timing (Went, 1949; Levine et al., 2008; Kimball et al., 2010; Walck et al., 2011; Mayfield et al., 2014; Huang et al., 2016). Climate change is already altering germination conditions through increasing temperatures and shifting precipitation regimes, leading to potential mismatches between formerly adaptive germination cues and both current and future environmental conditions (McNamara et al., 2011; Parmesan and Hanley, 2015; Bernhardt et al., 2020). Thus, shifts in climate may affect the timing of germination, the proportion of viable seeds that germinate (germination fraction), and the speed at which germination proceeds (germination rate). The degree to which the germination niche is constrained by evolutionary history could, in part, determine how species respond to shifting climate, with implications for individual fitness and population dynamics (Arène et al., 2017; Fang et al., 2017; Fernández-Pascual et al., 2021).

Germination timing determines the abiotic and biotic environments seedlings will experience, influencing performance during one of the most vulnerable plant life stages (Donohue et al., 2010; Matías et al., 2011; Gremer et al., 2016). To time germination with favorable conditions, germination cues may evolve such that seeds respond to partially or wholly reliable cues signaling favorable conditions for germination and establishment (Cohen, 1967, Donohue et al., 2010, Gremer et al., 2016; Bonamour et al., 2019). For example, a global-scale meta-analysis of 661 alpine species revealed their strong propensity to germinate in response to warm, wet conditions (Fernández-Pascual et al., 2021), which signal the onset of the summer growing season. On the other hand, in variable and changing environments, seeds may not germinate either because appropriate conditions or cues are unavailable, or because seeds may remain dormant even in the presence of those cues. Seed dormancy can act to distribute germination through time, spreading the risk of germinating into unfavorable conditions and acting as a bet hedging strategy (Cohen, 1966; Hoyle et al., 2015). Bet-hedging through germination is often found in unpredictable environments like deserts, where environmental cues are unreliable (Venable, 2007; Gremer and Venable, 2014). Thus, we may expect seeds from more variable environments to be less sensitive to seasonal cues and conditions and have lower germination, while those from environments with reliable cues will be more responsive to seasonal conditions. If seeds fail to germinate because of the absence of appropriate cues and conditions or due to seed dormancy, they must survive in the soil seedbank in order to germinate in future conditions.

Understanding variation among species’ germination strategies and responses to environmental cues for germination can reveal vulnerability or resilience to climate change (Kimball et al., 2010; Liu et al., 2013). Climate change is already altering both temperature and precipitation patterns and forecasts indicate continued change in coming decades (Wright et al., 2016; Pathak et al., 2018; Swain et al., 2018). For example, the IPCC (2021) reports global increases in temperature, shifts in the timing of precipitation, and increased variability in both temperature and precipitation, along with more frequent and intense extreme climatic events. In addition to rising temperatures, climate change has also shifted the timing of germination-triggering rains (Kimball et al., 2010; Levine et al., 2011; Mayfield et al., 2014; Pathak et al., 2018; Luković et al., 2021) both of which will affect germination conditions. For species with winter growing seasons, rainfall events arriving later in the season may occur under cooler temperatures, affecting germination fractions and rates (Kimball et al., 2010, 2011; Huang et al., 2016). In general, germination is expected to be faster and in higher fractions under warmer temperatures until a threshold is reached (Alvarado and Bradford 2002; Finch-Savage and Leubner-Metzger 2006), unless species have cold cued germination. Moreover, climate change may create mismatches between germination cues and conditions, such that seeds may germinate in unfavorable conditions or not germinate at all if they do not receive appropriate environmental cues (Donohue et al., 2010; Walck et al., 2011). Thus, climate change induced disruptions to both temperature and precipitation have implications for germination processes.

The *Streptanthus* (*s.l.*) clade of Brassicaceae are an ideal system to ask how shifting rainfall onset will alter germination patterns. The clade has desert origins and diversified as it moved northward (Cacho et al., 2021), as did other taxa comprising the Madro-tertiary geoflora (Axelrod, 1958). *Streptanthus* and *Caulanthus* species have also diversified across a range of typically drier environments in the Mediterranean climate of California (Cacho and Strauss, 2014; Cacho et al., 2021), characterized by cool wet winters and dry hot summers. Prior work in this system has shown that as these species spread from the desert, many of their traits diversified along the phylogeny, with closer relatives having more similar traits (Cacho and Strauss, 2014; Christie and Strauss, 2018). For example, Pearse et al. (2020) found phylogenetically conserved fitness responses to water availability, suggesting that diversification of those responses was constrained by evolutionary history. However, climate change has already shifted patterns of seasonal precipitation in Mediterranean climates (Giorgi and Lionello, 2008; Walck et al., 2011; Barredo et al., 2018). In California, the timing of first seasonal rains has shifted to later in the fall and winter growing season, resulting in less fall precipitation and more concentrated precipitation during the colder months (Luković et al., 2021); however, little is known about how these shifts influence germination rates and fractions.

To investigate how the timing of rainfall affects germination fraction and rate for *Streptanthus* and *Caulanthus* species across the *Streptanthus* (*s.l.*) clade, we experimentally varied the date of rainfall onset for 11 species and asked three questions, 1) How does variation in the timing of germination-triggering precipitation and corresponding seasonal temperature affect germination fraction and rate?, 2) What are the possible effects of contemporary shifts in precipitation timing on germination fraction and rate of species across the clade?, and 3) How have germination responses to seasonal conditions diversified across these closely related species? We expected that seeds would germinate at higher fractions and faster rates with earlier rainfall events that occur during warmer fall temperatures, which can facilitate germination as well as maximizing growing season length. We also expected that species adapted to divergent climates of origin would vary in germination response to rainfall onset timing. Specifically, we predicted that *Caulanthus* species that occupy drier, more variable environments, would be less affected by the timing of germination-triggering rains and corresponding temperatures than *Streptanthus* species, adapted to wetter, less variable environments (Golodets et al., 2013; Figure 1). Correspondingly, we also expected to see lower germination fractions, suggesting higher dormancy, in *Caulanthus* species from variable and less predictable local environments.

**Figure 1.**
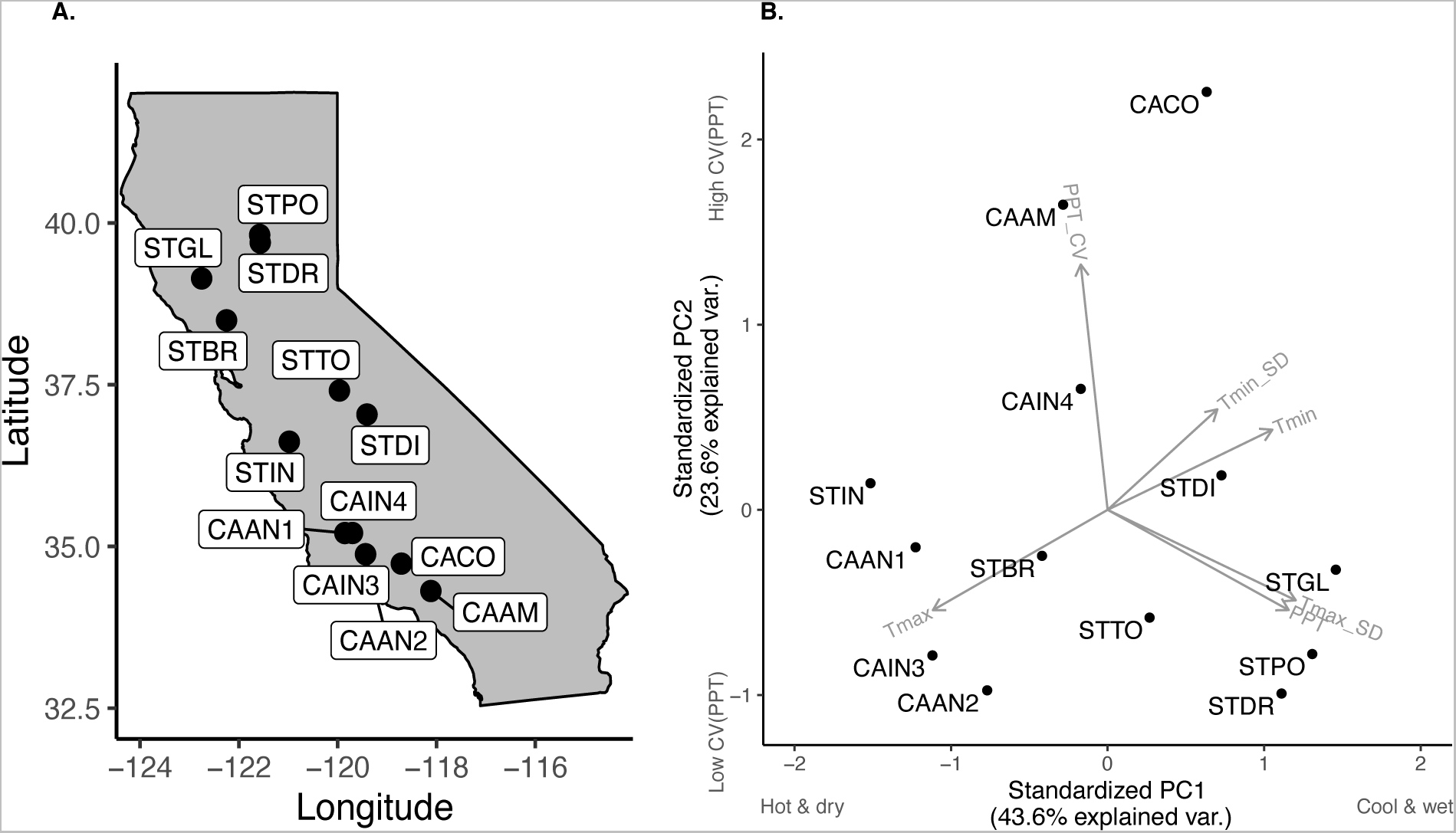
Location and climate for study species. (A) Seeds were collected from 13 populations of 11 species across California. See Table 1 for species abbreviations. (B) Principal component analysis of average germination season (September - December) climate for 1991-2015 for each location. First two principal components (PCs) illustrated on the x and y axes (Appendix S1: Table S11). Climate variables include precipitation (PPT), minimum and maximum temperature (Tmin, Tmax, respectively), as well as variability in precipitation (coefficient of variation, PPT_CV) and temperature (standard deviation of Tmin and Tmax, Tmin_SD and Tmax_SD respectively). Data source was Flint and Flint (2014).

**Table 1.** Species (11) and populations (13) included in this study from the *Caulanthus* and *Streptanthus* genera of Brassicaceae. Average germination season (September – December) climate data from 1991 – 2015 are also reported for each species’ location: precipitation (PPT, mm), maximum temperature (Tmax, °C), and minimum temperature (Tmin, °C). Climate data sourced from Flint and Flint (2014).

| **Species** | **Abbreviation** | **PPT** | **Tmax** | **Tmin** |
| --- | --- | --- | --- | --- |
| *Caulanthus amplexicaulis* | CAAM | 39.90 | 21.44 | 8.76 |
| *Caulanthus anceps* | CAAN1 | 16.42 | 22.98 | 5.93 |
| *Caulanthus anceps* | CAAN2 | 19.00 | 21.92 | 4.92 |
| *Caulanthus coulteri* | CACO | 21.74 | 18.71 | 8.22 |
| *Caulanthus inflatus* | CAIN3 | 19.97 | 21.97 | 5.77 |
| *Caulanthus inflatus* | CAIN4 | 18.07 | 21.03 | 9.04 |
| *Streptanthus breweri* | STBR | 66.99 | 21.71 | 8.45 |
| *Streptanthus diversifolius* | STDI | 58.07 | 20.61 | 8.08 |
| *Streptanthus drepanoides* | STDR | 100.18 | 21.66 | 9.61 |
| *Streptanthus glandulosus* | STGL | 124.75 | 18.45 | 8.29 |
| *Streptanthus insignis* | STIN | 30.53 | 22.27 | 5.47 |
| *Streptanthus polygaloides* | STPO | 142.42 | 19.82 | 7.93 |
| *Streptanthus tortuosus* | STTO | 50.76 | 22.11 | 8.25 |

## Materials and Methods

### Study System

To investigate how climatic shifts may influence germination fractions and rates, we used 13 populations from 11 species spanning the *Streptanthus clade* (*s.l.*; Thelypodieae Brassicaceae), which includes nonmonophyletic genera *Streptanthus* and *Caulanthus* (Table 1; Cacho et al., 2014). As a group, these species span the latitudinal range of the California Floristic Province and typically inhabit relatively barren, dry substrates ranging from sandy deserts to rocky and serpentine outcrops (Figure 1A; Cacho and Strauss 2014). The clade occupies habitats with a range of mean temperature and precipitation, and a range of variability in these measures (Figure 1B, Table 1; Pearse et al 2021; Cacho et al 2021). All species in this study experience the Mediterranean climate in California, in which the winter growing season begins with germination-triggering rain events in the fall or early winter and ends with the onset of summer drought.

### Phylogeny Estimation

Full methods for phylogeny estimation can be found in Cacho et al. (2014). In brief, the phylogenetic hypothesis was generated using six single copy nuclear genes, three identified specifically for this group in combination with three traditionally used nuclear regions (phyA, ITS, PEPC), and two chloroplast regions (trnL, trnH-psbA). The hypothesis was based on Bayesian MCMC runs consisting of three 50-million-generation independent runs with sampling every 5000 generations. For the two species with two populations, additional populations were added to the phylogeny at the same node with the same edge length as the other population of the species using the phytools (Revell, 2012) and ape (Paradis and Schliep, 2019) packages in R programming language (R Core Team 2021).

### Experimental Design

To assess species differences in germination responses to variation in the timing of germination-triggering rains, we experimentally imposed seven rainfall onset events throughout the germination season. Seven distinct germination cohorts were created by simulating germination-triggering rain events every two weeks in 2020: September 17, October 2, October 16, October 30, November 13, November 27, and December 11, 2020. The timing of these events encompasses natural interannual variation in the arrival of rains, as well as shifts observed with contemporary climate change (Gremer et al., 2020, Luković et al., 2021). This experiment was conducted in a “screenhouse” with a clear plastic roof that allowed for controlled watering, but exposed seeds to seasonal temperature fluctuations and changes in ambient light.

Seeds were collected as maternal families from field locations in 2019 and pooled for this experiment (Appendix S1: Table S1). For nine of the species, seeds were collected from one population approximately centrally located within the species’ range-wide temperature and precipitation space (Appendix S1: Figure S1). For two species (*C. anceps* and *C. inflatus*), for which we had more seed, we included seeds from two sites (Appendix S1: Figure S1). We assessed each population’s location in its species’ climate space by assembling a database of collection locations from the Consortium of California Herbaria (CCH2 Portal, 2023). We then matched collection locations with temperature and precipitation data extracted from the California Basic Characterization Model (Flint and Flint 2014) to perform a principal component analysis (Appendix S1: Table S2) of average germination season (September - December) climate for 1991-2015 of all species, highlighting the location of the population(s) of interest in this study (Appendix S1: Figure S1).

For each cohort, individual seeds were sown into 107 mL cone-tainer pots (Stuewe and Sons SC7) filled with a mix of 2/3 UC Davis potting soil (1:1:1 parts sand, compost, peat moss with dolomite) and ⅓ coarse 16 grit sand. For each species, except *C. coulteri*, 16 seeds were sown in each of three replicate blocks for a total of 48 seeds sown per cohort. For *C. coulteri*, for which we had limited seed, four seeds were sown per replicate block, 12 total seeds sown per cohort. Pots with seeds were randomly assigned to one of three blocks, which were arranged across three screenhouse benches.

We simulated germination-triggering rain events every two weeks by bottom-watering pots to saturation before sowing, then intensely misting pots for one week after sowing (Appendix S1: Table S3). Soil moisture conditions following the germination-triggering rain events were maintained by lower levels of watering (Appendix S1: Table S3). This procedure was repeated for each cohort with watering amounts following germination-triggering rain events reduced as the experiment progressed because of cooler seasonal temperatures. One week after the last cohort was sown, all cones were subjected to maintenance watering until germination surveys ceased.

Germination surveys were conducted daily in which every cone was censused and the germination date for each individual was recorded. By January 11, 2021, germination had slowed, and germination surveys were reduced to twice weekly and then further reduced on February 1, 2021, to once per week. All surveys and watering were completely stopped on April 16, 2021, after no germination had been observed in the previous two weeks.

To understand whether ungerminated seeds persisted in the soil as viable over the dry, Mediterranean California summer, all cones with ungerminated seeds were allowed to dry out and kept in the screenhouse to expose seeds to natural temperatures over the summer. The following fall, these pots were re-randomized, bottom-watered, and then received one week of high-frequency watering, as in the previous year, starting on September 15, 2021, to simulate the beginning of the next growing season (year 2). Germination surveys were then conducted daily until November 5, 2021, when they were reduced to three-times weekly. On November 22, 2021, surveys decreased to once per week and were stopped on December 27, 2021, after a two-week period without any germination.

### Thermal Germination Conditions

We determined the temperature each seed experienced prior to germination by calculating the mean temperature between the day of rainfall onset and day of germination from Thermochron DS1921G iButtons buried in soil-filled cones (Figure 2). These values were then used in models of germination rate. In models of germination fraction, the average was taken of the mean temperatures experienced by each seed in each block, within each cohort. This procedure gave a block-level estimate of mean temperature experienced by seeds in each block between rainfall onset date and germination date. Throughout the experiment, ambient temperatures decreased through time such that seeds in earlier rainfall onset events experienced warmer temperatures than seeds in later rainfall onset events (Figure 3A).

**Figure 2.**
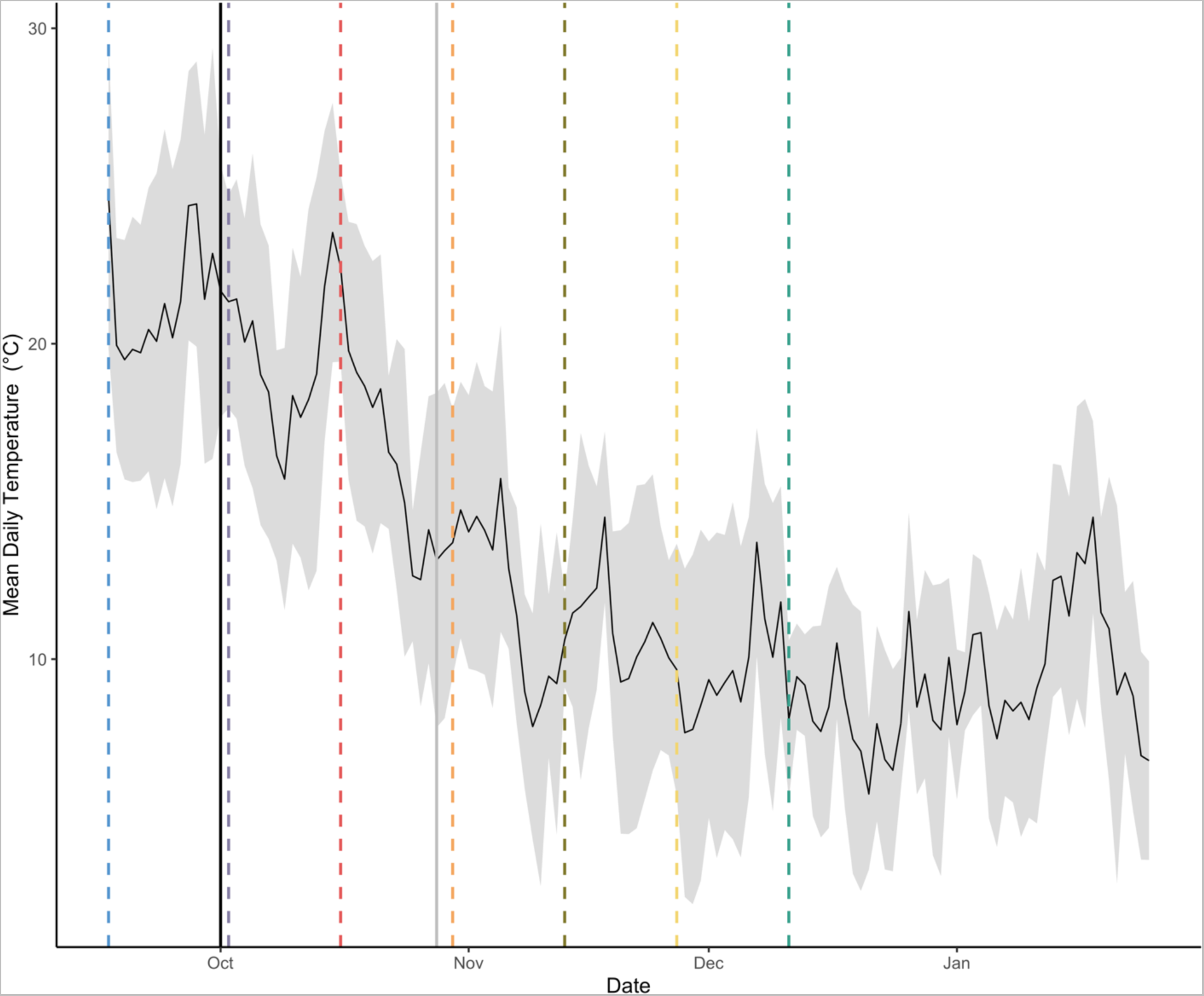
Mean and standard deviation of daily temperatures (°C) from the first rainfall onset date (September 17, 2020) to the last day of observed germination (January 25, 2021) in year one of the experiment. Dashed lines represent each rainfall onset date for each of seven cohorts with colors corresponding to those in Figure 3A. The solid black line corresponds to the date of historic onset of rainfall (October 1) and the solid gray line represents the 27 day shift to the contemporary date of rainfall onset (October 28) in California according to Luković et al. (2021).

**Figure 3.**
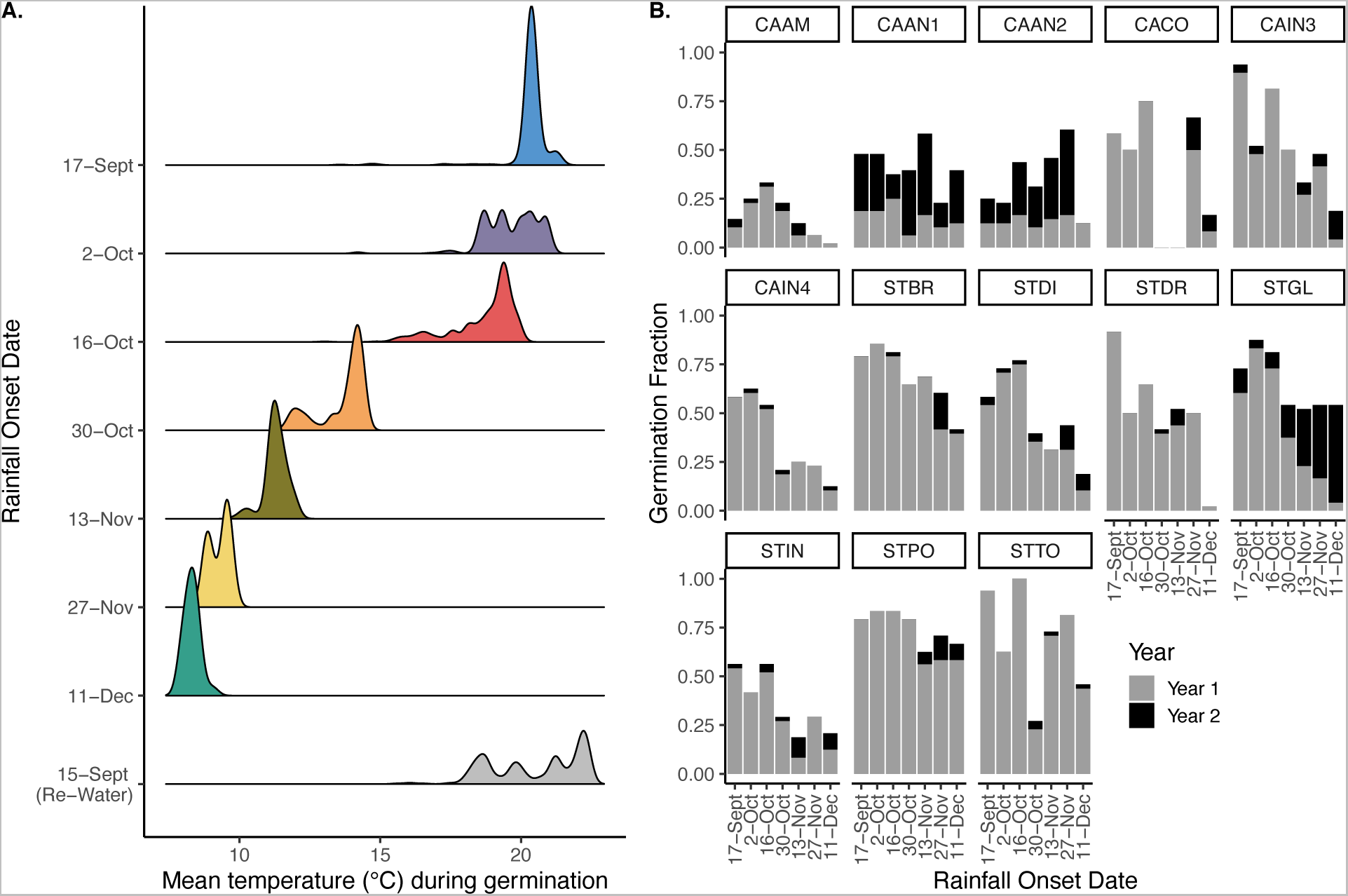
Germination fractions and temperatures seeds experienced during the study. (A) Distribution of mean temperature seeds in each cohort experienced between their rainfall onset date, displayed on the y-axis, and germination date. Seeds that did not germinate during year one of the experiment (September 17, 2020, to April 16, 2021) were re-watered on September 15, 2021, and germination was followed until December 6, 2021. (B) Amount of total germination fraction for each species during year one versus year two of the study at each rainfall onset date.

### Data Analysis

#### Germination Fraction and Rate

Germination fraction was calculated at the block level as the number of germinated seeds divided by the total number of seeds sown. Germination rate was calculated for each seed as the reciprocal of the number of days between germination date and rainfall onset date (Bewley et al., 2013). Grouped logistic regressions were used to evaluate relationships between germination fraction and seasonal temperature. Regressions were built with groups comprising the seeds sown for each block (3), and successes indicating germination, with block included as a random effect. For germination rate, these relationships were evaluated using linear mixed effects models (Bates et al., 2015) with block as a random effect. Relationships between germination fraction, rate, and the timing of rainfall onset were evaluated for each species separately. Both linear and quadratic models were fit, and likelihood ratio tests were used to determine the significance of factors in the models. Only linear model results are presented as their results were not distinguishable from quadratic models. Marginal means were estimated for each species using the emmeans package (Lenth, 2022) for use in *post hoc* comparisons.

We also tested whether species differ in their germination responsiveness to the timing of rainfall onset. A Pearson’s Chi-squared test with three contingencies (Year 1, Year 2, Never) was used to evaluate germination timing. We performed *post hoc* analyses using the chisq.posthoc.test package (Ebbert, 2019) in R with a Bonferroni p-value adjustment.

#### Potential Effects of Climate Change

To understand how climate change-induced shifts in precipitation timing affect germination responses, we compared germination responses (fractions and rates) on dates that represent mean historical and contemporary rainfall onset based on Luković et al. (2021), who showed that since 1960, rainfall onset date has shifted approximately 27 days later into the fall from an average rainfall onset date of October 1 (1960-1989) to an average onset date of October 28 (1990-2019). These dates align closely with our rainfall onset dates of October 2 (historical date) and October 30 (contemporary date). Thus, we compared germination fractions and rates of historical and contemporary rainfall onset dates using species-specific generalized linear models with binomial error and a logit link function for germination fraction and linear models for germination rate. Marginal means were estimated, and comparisons between historical and contemporary rainfall onset dates were made using the emmeans package (Lenth, 2022) with p-values adjusted using the Tukey method for multiple comparisons.

#### Phylogenetic Analyses of Species’ Responses

We compared germination responses to rainfall onset date and seasonal temperatures across the phylogeny by extracting the slopes from the relationships between germination fraction and rate in response to rainfall onset and temperature for each species and testing for phylogenetic signals in slopes. Estimates of phylogenetic signals allow us to understand how these responses have evolved as these species spread and diversified into different environments, and whether germination behavior is constrained by evolutionary history. Using the R package phytools (Revell, 2012), we estimated phylogenetic signals using Pagel’s **λ**. We tested the null hypothesis that Pagel’s **λ** = 0, where a value equal to 0 indicates low phylogenetic signal, evolution of germination responses is independent of phylogeny, and a value equal to 1 indicates high phylogenetic signal, differences in species’ responses are proportional to their shared history (Pagel, 1999). Intermediate values of Pagel’s **λ** suggest phylogenetic relatedness is not the only factor contributing to the observed patterns.

## Results

In general, later rainfall onset had a negative effect on both germination fraction (Figure 4, Appendix S1: Table S4) and germination rate (Figure 5, Appendix S1: Table S5), though responses varied among species. Germination fraction significantly decreased at cooler temperatures for all but three species (*C. amplexicaulis*, *C. anceps*, and *C. coulteri*) while germination rate significantly decreased for all species (Figures 6-7; Appendix S1: Table S6-S7).

**Figure 4.**
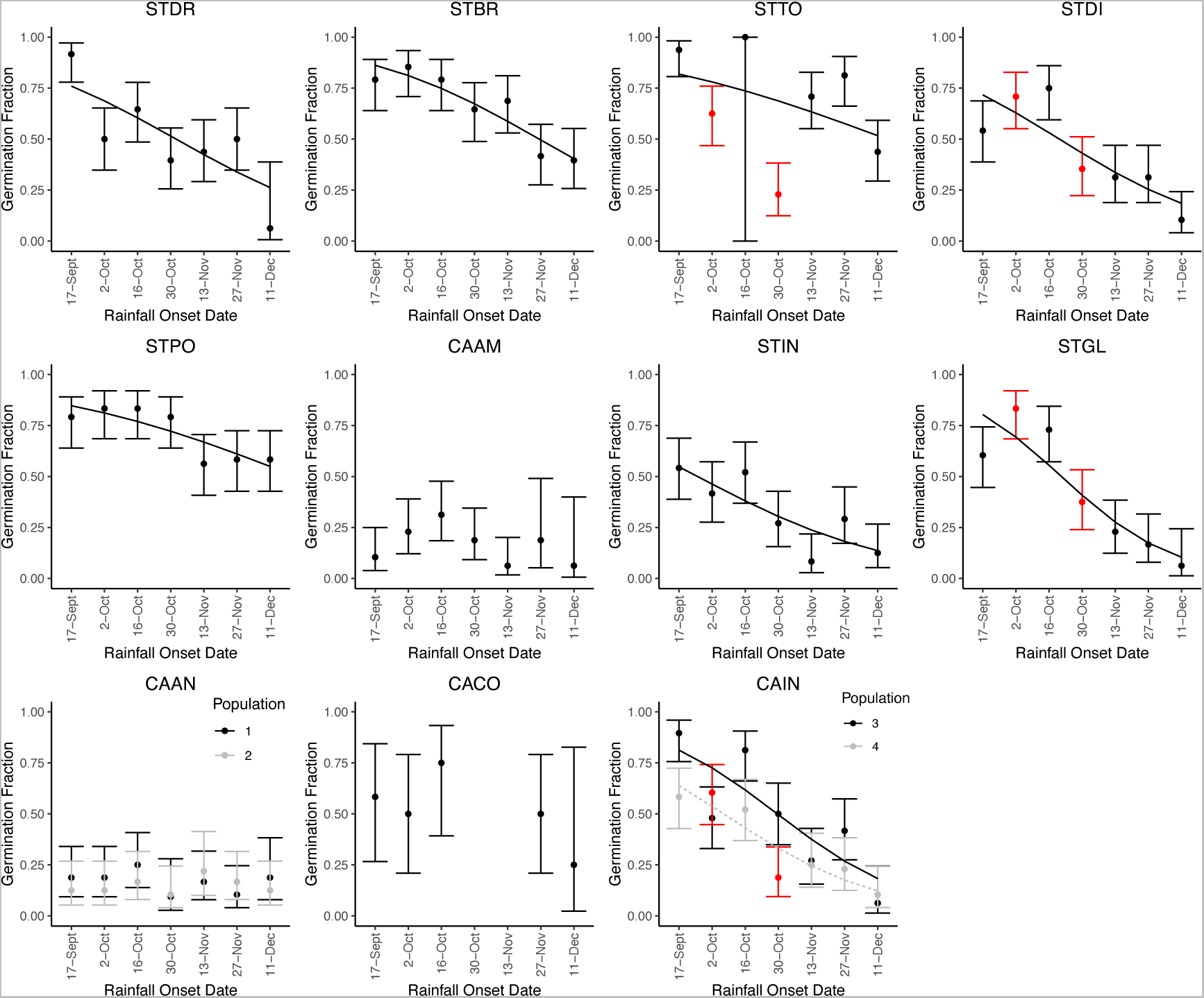
Relationships between germination fraction and rainfall onset date and comparison of germination fraction between 2-Oct (historical onset of rainfall) and 30-Oct (contemporary onset of rainfall). Plots where regression lines are present had significant relationships between germination fraction and rainfall onset date (back transformed from logit scale). For all significant relationships, germination fraction decreased as rainfall onset occurred later in the season. Points represent germination fraction for each cohort (mean ± 1 SE, back transformed from logit scale). In plots where points and SE bars are red, a significant decrease in germination fraction was found between the historical onset date of rainfall (2-Oct) and the contemporary onset date of rainfall (30-Oct).

**Figure 5.**
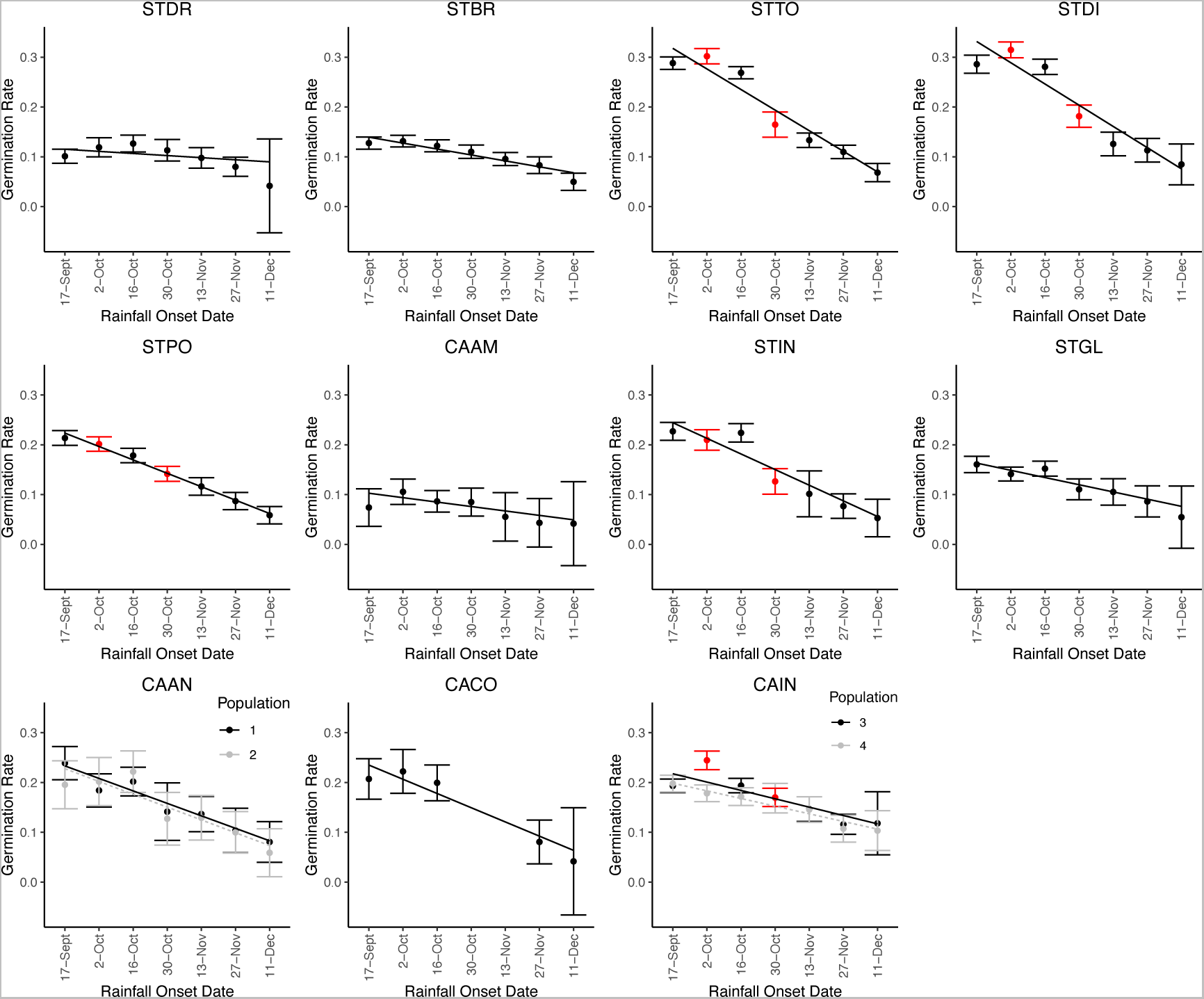
Relationships between germination rate (1/days to germination) and rainfall onset date and comparison of germination rate between 2-Oct (historical onset of rainfall) and 30-Oct (contemporary onset of rainfall). Regression lines represent significant, negative relationships between germination rate and rainfall onset date, where germination rate decreased as rainfall onset occurred later in the season. Points represent germination rate for each cohort (mean ± 1 SE). In plots where points and SE bars are red, a significant decrease in germination rate was found between the historical onset date of rainfall (2-Oct) and the contemporary onset date of rainfall (30-Oct).

**Figure 6.**
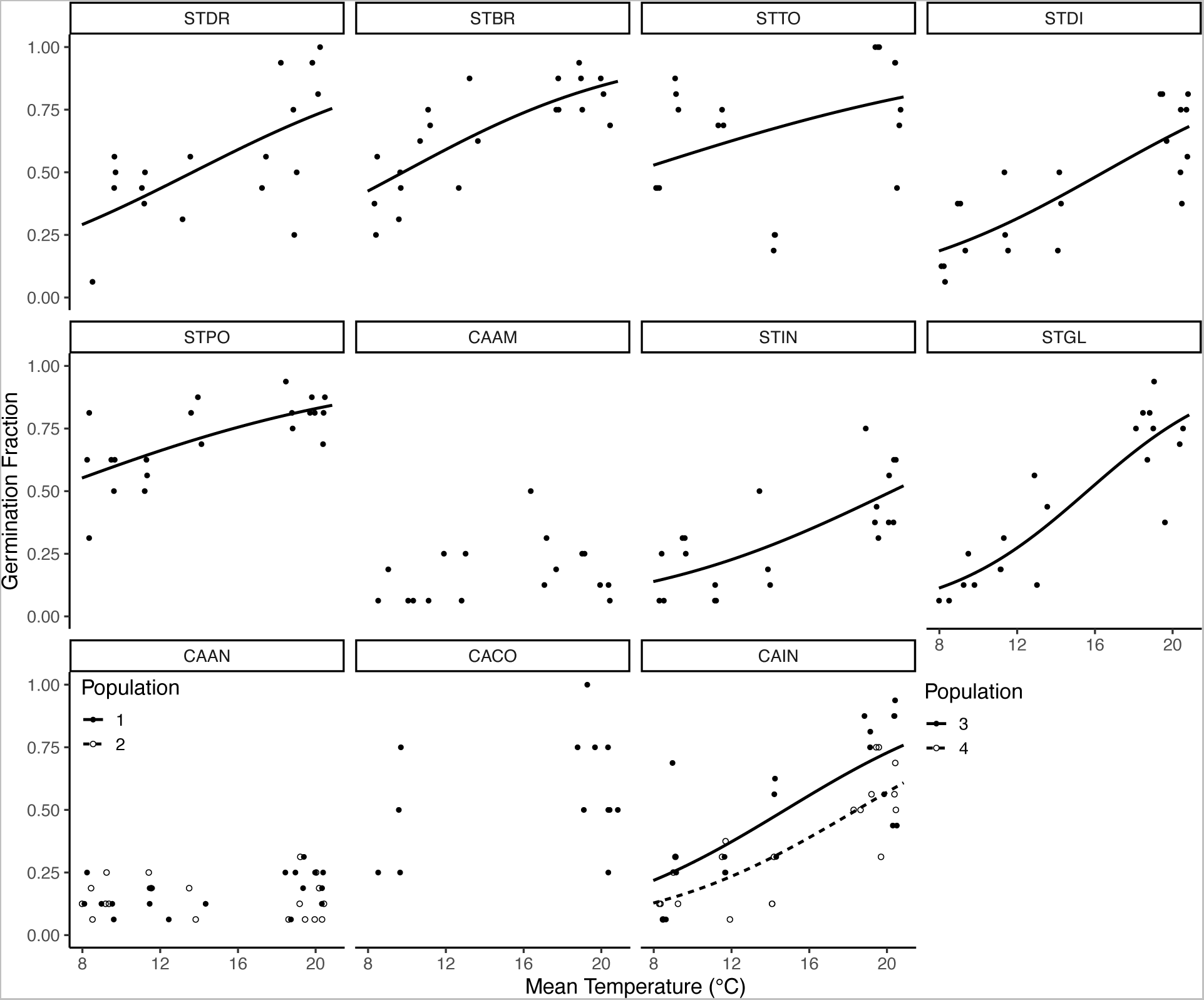
Relationships between germination fraction and mean temperature seeds experienced between their rainfall onset date and germination date. Regression lines represent significant, positive relationships between germination fraction and mean temperature seeds experienced (back transformed from logit scale).

**Figure 7.**
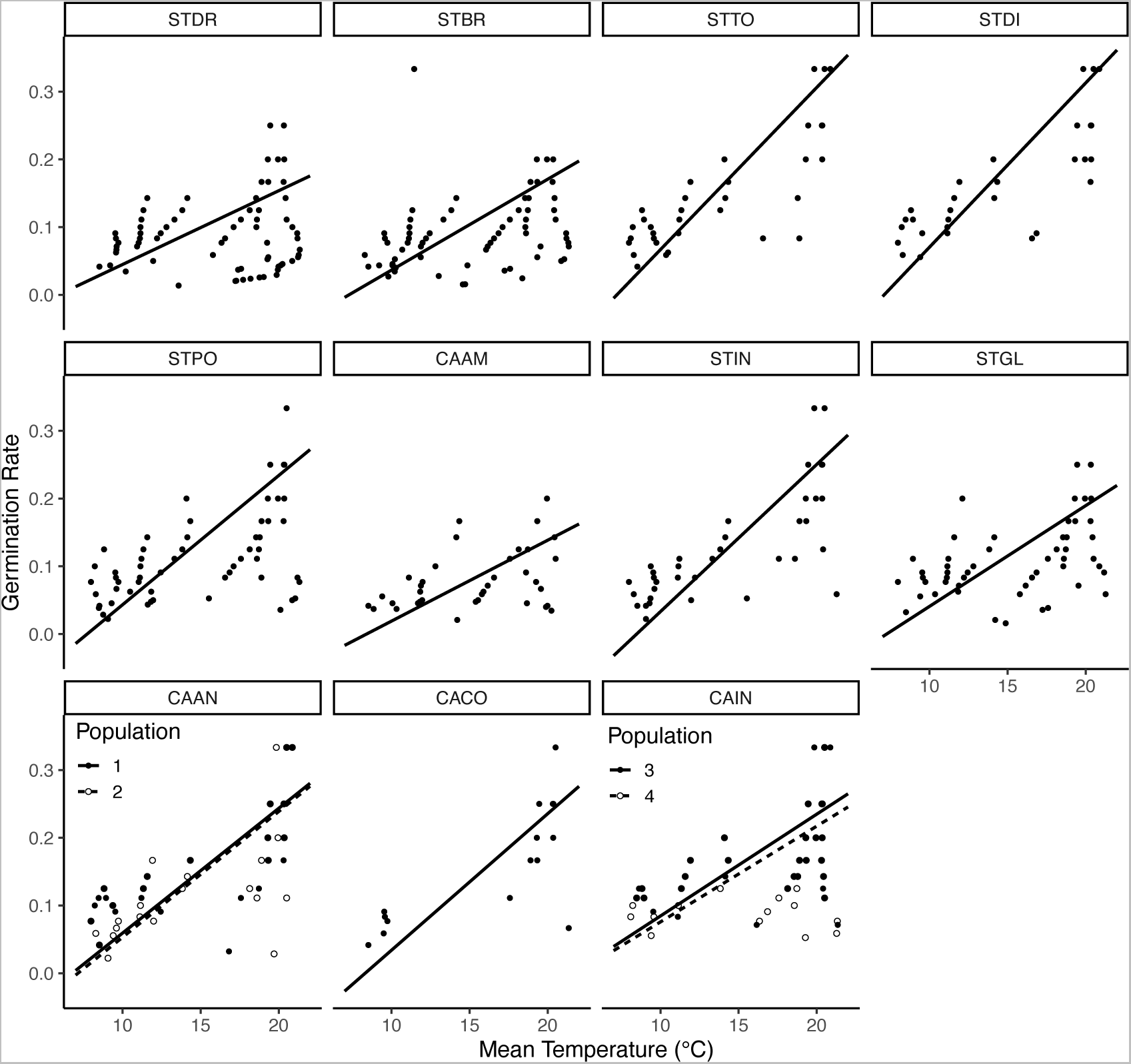
Relationships between germination rate and mean temperature seeds experienced between their rainfall onset date and germination date. Regression lines represent significant, positive relationships between germination rate (1/days to germination) and mean temperature seeds experienced.

To understand how climate change-induced shifts in precipitation timing may affect germination responses, we compared historical and contemporary timings of rainfall onset. Germination fraction significantly decreased between historical (cohort 2) and contemporary (cohort 4) rainfall timing for *S. diversifolius*, *S. glandulosus*, *S. tortuosus*, and one of the populations of *C. inflatus* (CAIN4; Figure 4; Appendix S1: Table S8). Germination rate significantly decreased for *S. diversifolius*, *S. insignis*, *S. polygaloides*, *S. tortuosus*, and the other population of *C. inflatus* (CAIN3; Figure 5; Appendix S1: Table S9).

Slopes of relationships between germination rate and the timing of rainfall onset (**λ** = 0.99, P < 0.005) and between germination rate and the mean temperature seeds experienced (**λ** = 0.99, P < 0.05) showed significant phylogenetic signal (Figure 8). Slopes of relationships between germination fraction, the timing of rainfall onset (**λ** = 0.87, P = 0.05), and the mean temperature seeds experienced (**λ** = 0.84, P = 0.08) showed moderate, but non-significant phylogenetic signal.

**Figure 8.**
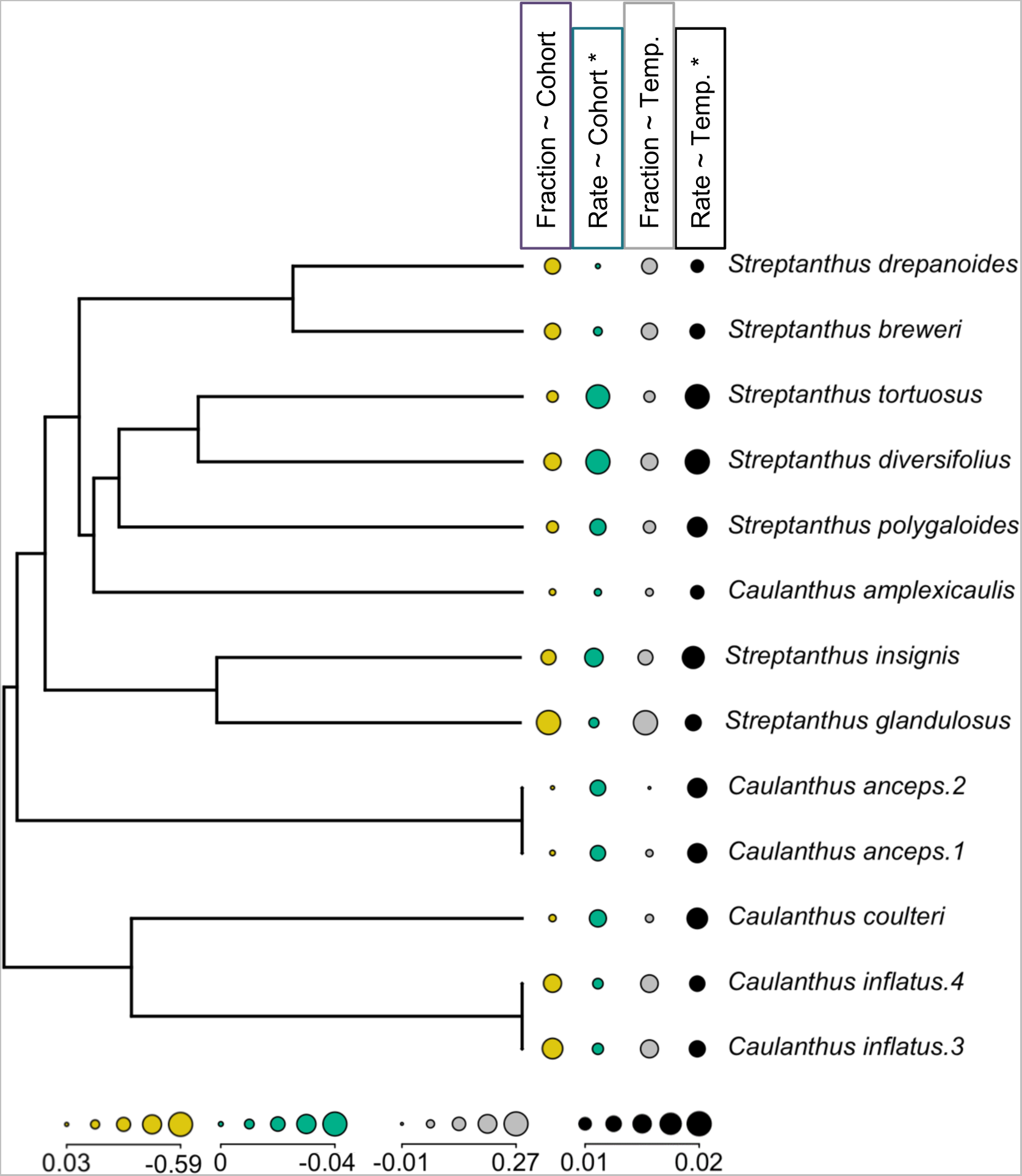
Slopes of the relationships evaluated in this study displayed across the phylogeny. Yellow dots display the slopes of the relationship between germination fraction and rainfall onset date (Fraction ∼ Cohort) (Figure 4). All slopes are negative except for *Caulanthus anceps.2* (0.03). Green dots display the slopes of the negative relationships between germination rate and rainfall onset date (Rate ∼ Cohort) (Figure 5). Gray dots display the slopes of the relationship between germination fraction and mean temperature experienced by seeds (Figure 6). All slopes are positive except for *Caulanthus anceps.2* (-0.01). Black dots display the positive slopes of the relationships between germination rate and mean temperature experienced by seeds (Figure 7). Relationships with an * showed significant phylogenetic signal according to Pagel’s **λ**.

Species differed in the fates of their seeds across the study (germination in year 1, fall year 2, or never; chi-squared = 1006.1, df = 24, p < 0.0001; Figure 3A). While most species germinated much less in the second year, three species had different responses (Figure 3A; Appendix S1: Table S10). However, both populations of *C. anceps*, a desert dwelling species, had higher germination in year 2 than in year 1. *S. glandulosus* had low germination in the later rainfall onset events of year 1, which then germinated in high fractions in the early season rainfall event of year 2 (Figure 3A: 27-Nov. 11-Dec.). *C. amplexicaulis*, a mid-high elevation lower latitude species, was not as sensitive to the timing of rainfall onset or temperature, with significantly lower germination in both years (Figure 3A; Appendix S1: Table S10).

## Discussion

Here, we assessed how variation in the timing of the onset of rain events affected germination fraction and rate for closely related species that have radiated into different geographic regions within Mediterranean climates of California. Our results indicate that later onset of seasonal rains during cooler seasonal temperatures generally decreases germination fractions and rates, though species varied in their sensitivity of responses. Further, comparisons across cohorts indicate that six species may already be experiencing negative effects of contemporary shifts in the timing of rainfall. Species differed in responsiveness to seasonal cues affecting germination timing: *Caulanthus* species from drier (*C. anceps*) and more variable (*C. amplexicaulis* and *C. coulteri*) sites were less sensitive to the timing of rainfall onset events or their associated temperatures than species from wetter, cooler environments. Variation in responses to the timing of rainfall and corresponding temperatures across the clade show phylogenetic signal, which may indicate that diversification of germination responses was constrained by evolutionary history as species radiated into different climates.

In systems with winter growing seasons, like lower elevation habitats of California, earlier onset of seasonal rains typically corresponds with warmer temperatures and may also lead to a longer growing season. Thus, earlier rains may drive higher and faster germination simply due to thermal requirements for germination or may be an adaptive response in which seeds are tuned to early conditions for germination in order to take advantage of a longer growing season. On the other hand, there may be advantages to germinating later in the season during cooler temperatures that may indicate that reliable cool, wet weather has arrived (Mayfield et al., 2014). In the current study, germination rate significantly decreased for all species with later onset of precipitation that occurred under cooler temperatures. Germination fraction also showed this pattern for most species. Huang et al. (2016) reported similar results in a Sonoran Desert plant community finding longer thermal times to germination (slower germination rates) and lower germination percentages later in the season under cooler conditions. In their system and in ours, most of the variability in growing season length comes from the onset of germination-triggering rains, which start the growing season, instead of the timing of the onset of dry conditions that end it (Kimball et al., 2011; Luković et al., 2021). Thus, earlier germination, under warmer temperatures, is the main avenue to increase growing season length. Together, our findings and previous research highlight how important the interaction of precipitation timing and temperature is for driving germination patterns and suggest that shifting rainfall timing could overwhelm effects of increasing temperatures with climate change.

Indeed, a growing body of literature has found that germination cues interact to drive climate change effects on germination timing, fractions, and rates (Levine et al., 2008; Donohue et al., 2010; Kimbell et al., 2010; Levine et al., 2011; Huang et al., 2016). While the timing of precipitation onset has shifted later since 1960 in California (Luković et al., 2021), average temperature has not followed the same trend as sharply as precipitation (Rapacciuolo et al., 2014; Wright et al., 2016; Pathak et al., 2018), suggesting a recent and continuing mismatch between the timing of precipitation and the temperature following rain events relative to historic patterns. Here, we found that six species in this study showed decreases in germination fractions or rates between historic and contemporary onsets of precipitation, though two species, *S. diversifolius* and *S. tortuosus* showed significant decreases in both, suggesting they may already be experiencing more extreme effects of shifting precipitation timing than other species. This result highlights species-specific sensitivities to climate change, calling for species-level restoration and conservation plans (Barga et al., 2017; Finch-Savage and Footitt, 2017).

Species in this study showed variation in relationships between germination responses and environmental cues for germination, and phylogenetic signal suggests that the adaptive diversification of these responses may have been constrained by evolutionary history as these species diverged. Previous work has also found strong phylogenetic signal in germination cues (Arène et al., 2017; Fernández-Pascual et al., 2021; Baskin et al., 2022). In the *Streptanthus* (*s.l.*) clade, research has linked divergence and spread of these species to edaphic and climatic adaptations, including soil nutrients and precipitation quantity (Cacho and Strauss, 2014; Christie and Strauss, 2018; Pearse et al., 2020). In this study, we found that many of the *Caulanthus* species are much less sensitive to rainfall timing and corresponding temperatures than *Streptanthus* species. Lower latitude *Caulanthus* species have typically evolved in drier, warmer, and more variable habitats than higher latitude *Streptanthus* species (Christie and Strauss, 2018), potentially contributing to divergence in their germination sensitivities. Differences in sensitivity of species to environmental cues for germination has implications for the success of germination and population dynamics in future climate scenarios. With continued climate change, seeds may either not germinate or germinate under suboptimal conditions based on formerly adaptive germination cues, unless they are able to shift germination timing by tracking conditions through time or space (Catelotti et al., 2020), adapt to respond to new combinations of germination cues (Donohue et al., 2010; Walck et al., 2011), or use a combination of these strategies (Gremer et al., 2016; Gremer et al., 2020a).

Germination strategies can include using cues to time germination with appropriate conditions, not germinating in a given year if germination conditions or cues are unavailable, or remaining dormant in the soil. Species in this study encompassed all these germination strategies. First, most species were quite responsive to the interaction of precipitation and temperature germination cues. On the other hand, *S. glandulosus* seeds did not germinate at all in later cohorts, instead waiting until the following fall to germinate in warmer temperatures (Figure 3A), indicating that appropriate cues were not available in later cohorts of the first year. Such patterns can be achieved through secondary dormancy, which can prevent germination under unfavorable conditions and must be released under favorable conditions in subsequent years (Probert, 2000; Walck et al., 2011; Finch-Savage and Footitt, 2017; Hawkins et al. 2017). In contrast, three *Caulanthus* species had similar low germination fractions regardless of the timing of rainfall onset or corresponding temperatures. Low germination across conditions is consistent with bet-hedging strategies to spread germination across years, particularly in harsh and temporally variable environmental conditions (Cohen, 1966; Venable, 2007; Gremer and Venable, 2014). Together, these patterns suggest a range of germination strategies have evolved along the phylogeny.

Overall, our findings shed further light on germination sensitivity to changing temperature and precipitation and suggest responses have evolved as the clade diversified into different climatic conditions. While populations were chosen for their approximate central location within climate space for their species (Appendix S1: Figure S1), as well as availability of seed from naturally occurring populations, a caveat is that these represent one to two populations per species. Indeed, our study trades-off replicating populations within species for comparing species across the clade, however we can still infer some species-level properties. For instance, a population of *C. anceps* (CAAN2) and a population of *C. inflatus* (CAIN3) were collected from the same field location (Appendix S1: Table S1), but each of these populations reacts more similarity to the other population of its species than to the population of the other species from the same environment (Figures 4-7). Moreover, one would be unlikely to detect phylogenetic signal in germination properties if intra-and interspecific variation were similar (see also Pearse et al 2021 for water use evolution in this clade). Another caveat is that we focused on temperature and precipitation since these cues and their interaction has already shifted with contemporary climate change. However, other seasonal cues may influence responses, such as photoperiod, chilling, or fire (Baskin and Baskin, 2014), and these cues are also likely shifting with anthropogenic change (IPCC 2021).

Together, our results highlight how climate change is progressing faster than germination niche evolution in some species, creating mismatches between formerly adaptive germination cues and optimal seasonal conditions. This trend has implications for population persistence and species dynamics (Donohue et al., 2010; Huang et al., 2016; Bernhardt et al., 2020; Kimball et al. 2010, 2011, Levine et al. 2011, Gremer et al. 2020a). Our results underscore the importance of temperature in mismatched responses and allowed us to identify which species may be more sensitive to current and future shifts associated with climate change. The next critical steps are to link these responses with fitness and population dynamics, using controlled experiments, demographic data, and process-based models that can predict germination timing in response to current and future climate (Alvardo and Bradford 2002; Huang et al. 2016; Hamann et al., 2021). These directions are vital to further our understanding of how climate factors drive germination and scale to affect population persistence and species distributions.

## Acknowledgements

The authors thank Natascha Paxton and Shannon Reilly for assistance gathering data. Funding was provided by NSF DEB (1831913 to J.R. Gremer, J. Schmitt, S.Y. Strauss, and J. Maloof.

## Authors’ Contributions

J.R.G, J.S., S.Y.S. and J.N.M conceived and designed the study. A.M, E.C., S.R.A, J.S performed the experiment, with help from J.R.G. and S.Y.S in data collection. S.J.W., J.R.G., S.Y.S., and J.S designed the data analyses. S.J.W. and J.R.G. analyzed the data with assistance from J.S., S.Y.S, A.M., S.R.A., and E.C. S.J.W., J.R.G. A.M., and J.S. wrote the manuscript with contributions from all other authors. All authors contributed to development of ideas, analyses, and interpretation of results.

