## Supplementary Material for "Germination responses to changing rainfall timing reveal potential climate vulnerability in a clade of wildflowers"

**Table S1.** Species (11) and populations (13) included in this study from the *Caulanthus* and *Streptanthus* genera of Brassicaceae. Full population names and location information are given.

| **Species** | **Abbreviation** | **Population** | **Latitude** | **Longitude** |
| --- | --- | --- | --- | --- |
| *Caulanthus amplexicaulis* | CAAM | Camp Colby | 34.30539 | -118.11469 |
| *Caulanthus*  *anceps* | CAAN1 | Carrizo Plain | 35.20738 | -119.8539 |
| *Caulanthus*  *anceps* | CAAN2 | Ballinger Canyon | 34.87855 | -119.43405 |
| *Caulanthus*  *coulteri* | CACO1 | Frazer Road | 34.73461 | -118.71113 |
| *Caulanthus*  *inflatus* | CAIN3 | Ballinger Canyon | 34.88316 | -119.46696 |
| *Caulanthus*  *inflatus* | CAIN4 | Hurricane Road | 35.20804 | -119.69993 |
| *Streptanthus*  *breweri* | STBR3 | Turtle Rock | 38.496115 | -122.2485 |
| *Streptanthus diversifolius* | STDI | Big Sandy | 37.04 | -119.4068 |
| *Streptanthus drepanoides* | STDR2 | Lime Saddle | 39.698016 | -121.56693 |
| *Streptanthus glandulosus* | STGL1 | Bartlett Springs | 39.14296 | -122.75256 |
| *Streptanthus*  *insignis* | STIN | Panoche Road | 36.61968 | -120.97869 |
| *Streptanthus polygaloides* | STPO1 | Magalia | 39.815153 | -121.57857 |
| *Streptanthus*  *tortuosus* | STTO-BH | Ben Hur | 37.40985 | -119.96458 |

**Table S2.** Principal component analysis of average germination season (September - December) climate for 1991-2015 for species’ locations taken from herbarium records from the Consortium of California Herbaria. The first two principal component axes (PCs) are illustrated for each species in Figure S1. Climate variables include precipitation (PPT), minimum and maximum temperature (Tmin, Tmax, respectively), as well as variability in precipitation (coefficient of variation, PPT_CV) and temperature (standard deviation of Tmin and Tmax, Tmin_SD and Tmax_SD respectively). Data source was Flint and Flint, (2014).

| **Variable** | **PC1** | **PC2** |
| --- | --- | --- |
| PPT | 0.54 | -0.22 |
| PPT_CV | -0.42 | 0.25 |
| Tmin | -0.45 | -0.21 |
| Tmin_SD | -0.15 | -0.52 |
| Tmax | -0.55 | 0.001 |
| Tmax_SD | -0.07 | -0.76 |
| Cumulative Proportion | 0.39 | 0.61 |

**Table S3.** Dates and details on the watering regime for each of the seven cohorts in this study. Pots were bottom watered before sowing seeds to saturate the soil. This saturation was maintained by misting pots under the high frequency program for one week after sowing. Pots were watered under the low frequency program the following week to simulate field soil moisture conditions following large rain events. This process was repeated for each of the cohorts.

| **Cohort** | **Bottom Water Date** | **Planting Date** | **High Frequency Program** | **Low Frequency Program** |
| --- | --- | --- | --- | --- |
| Cohort 1 | 09/16/2020 | 09/17/2020 | 5 minutes, 7x per day | 5 minutes, 3x per day |
| Cohort 2 | 10/01/2020 | 10/02/2020 | 5 minutes, 7x per day | 5 minutes, 3x per day |
| Cohort 3 | 10/15/2020 | 10/16/2020 | 5 minutes, 7x per day | 5 minutes, 3x per day |
| Cohort 4 | 10/28/2020 | 10/30/2020 | 5 minutes, 7x per day | 5 minutes, 3x per day |
| Cohort 5 | 11/11/2020 | 11/13/2020 | 5 minutes, 6x per day | 5 minutes, 3x per day |
| Cohort 6 | 11/25/2020 | 11/27/2020 | 5 minutes, 2x per day | 5 minutes, 1x per day |
| Cohort 7 | 12/09/2020 | 12/11/2020 | 5 minutes, 2x per day | 5 minutes, 1x per day |

**Table S4.** Results from grouped logistic regressions with germination fraction as the response variable and rainfall onset date as the predictor variable. Coefficient estimates are reported for the Intercept and Rainfall Onset Date terms from the models along with the p-value associated with the Rainfall Onset Date term. P-values in bold are significant.

| **Species** | **Intercept** | **Rainfall Onset Date** | **P-value** |
| --- | --- | --- | --- |
| CAAM | -1.22 | -0.11 | 0.26 |
| CAAN1 | -1.29 | -0.08 | 0.33 |
| CAAN2 | -1.91 | 0.03 | 0.68 |
| CACO | 0.64 | -0.12 | 0.36 |
| CAIN3 | 1.96 | -0.49 | **< 0.0001** |
| CAIN4 | 0.99 | -0.42 | **< 0.0001** |
| STBR | 2.20 | -0.37 | **< 0.0001** |
| STDI | 1.33 | -0.40 | **< 0.0001** |
| STDR | 1.52 | -0.36 | **< 0.0001** |
| STGL | 2.00 | -0.59 | **< 0.0001** |
| STIN | 0.53 | -0.33 | **< 0.0001** |
| STPO | 1.96 | -0.25 | **< 0.0001** |
| STTO | 1.75 | -0.24 | **< 0.0001** |

**Table S5.** Results from linear regressions with germination rate as the response variable and rainfall onset date as the predictor variable. Coefficient estimates are reported for the Intercept and Rainfall Onset Date terms from the models along with the p-value associated with the Rainfall Onset Date term. Adjusted R^2^ values for the models are also given. P-values in bold are significant.

| **Species** | **Intercept** | **Rainfall Onset Date** | **P-value** | **R^2^** |
| --- | --- | --- | --- | --- |
| CAAM | 0.11 | -0.009 | **0.04** | 0.07 |
| CAAN1 | 0.26 | -0.06 | **< 0.0001** | 0.50 |
| CAAN2 | 0.25 | -0.03 | **< 0.0001** | 0.41 |
| CACO | 0.26 | -0.03 | **< 0.0001** | 0.50 |
| CAIN3 | 0.23 | -0.02 | **< 0.0001** | 0.25 |
| CAIN4 | 0.21 | -0.02 | **< 0.0001** | 0.27 |
| STBR | 0.15 | -0.01 | **< 0.0001** | 0.25 |
| STDI | 0.37 | -0.04 | **< 0.0001** | 0.63 |
| STDR | 0.12 | -0.004 | **< 0.0001** | 0.02 |
| STGL | 0.18 | -0.01 | **< 0.0001** | 0.18 |
| STIN | 0.28 | -0.03 | **< 0.0001** | 0.56 |
| STPO | 0.25 | -0.03 | **< 0.0001** | 0.56 |
| STTO | 0.36 | -0.04 | **< 0.0001** | 0.75 |

**Table S6.** Results from grouped logistic regressions with germination fraction as the response variable, mean temperature seeds experienced as the predictor variable, and block included as a random effect. Least-squares means were estimated for each species and are reported along with 95% confidence intervals (CI) used to evaluate significance. CI values in bold are significant.

| **Species** | **Temperature** | **CI** |
| --- | --- | --- |
| CAAM | 0.05 | -0.02 – 0.13 |
| CAAN1 | 0.05 | -0.02 – 0.11 |
| CAAN2 | -0.01 | -0.08 – 0.05 |
| CACO | 0.06 | -0.05 – 0.17 |
| CAIN3 | 0.19 | **0.14 – 0.24** |
| CAIN4 | 0.18 | **0.13 – 0.24** |
| STBR | 0.17 | **0.11 – 0.22** |
| STDI | 0.17 | **0.13 – 0.22** |
| STDR | 0.16 | **0.10 – 0.22** |
| STGL | 0.27 | **0.21 – 0.33** |
| STIN | 0.15 | **0.10 – 0.20** |
| STPO | 0.11 | **0.06 – 0.17** |
| STTO | 0.10 | **0.05 – 0.15** |

**Table S7.** Results from linear mixed-effects model with germination rates as the response variable, mean temperature seeds experienced, species, their interaction, and cohort as fixed effects, and block included as a random effect. Least-squares means were estimated for each species and are reported along with 95% confidence intervals (CI) used to evaluate significance. CI values in bold are significant.

| **Species** | **Temperature** | **CI** |
| --- | --- | --- |
| CAAM | 0.01 | **0.01 – 0.02** |
| CAAN1 | 0.02 | **0.02 – 0.02** |
| CAAN2 | 0.02 | **0.02 – 0.02** |
| CACO | 0.02 | **0.02 – 0.02** |
| CAIN3 | 0.02 | **0.01 – 0.02** |
| CAIN4 | 0.01 | **0.01 – 0.02** |
| STBR | 0.01 | **0.01 – 0.02** |
| STDI | 0.02 | **0.02 – 0.03** |
| STDR | 0.01 | **0.01 – 0.01** |
| STGL | 0.01 | **0.01 – 0.02** |
| STIN | 0.02 | **0.02 – 0.02** |
| STOP | 0.02 | **0.02 – 0.02** |
| STTO | 0.02 | **0.02 – 0.03** |

**Table S8.** Results from comparisons of germination fractions between historical (Oct-2) and contemporary (30-Oct) onset of seasonal precipitation. Grouped logistic regressions were fit with germination fraction as the response variable and the timing of rainfall onset (cohort) as the predictor variable. Comparisons of estimated marginal means between rainfall onset dates were made using the emmeans package (Lenth 2022). Least-squares means are reported for Oct-2 and Oct-30, their Contrast value, and Tukey adjusted p-values of the contrasts. P-values in bold are significant. No comparison was made for CACO due to lack of germination in the Oct-30 rainfall event.

| **Species** | **Oct-2** | **Oct-30** | **Contrast** | **P-value** |
| --- | --- | --- | --- | --- |
| CAAM | -1.21 | -1.47 | 0.25 | 0.99 |
| CAAN1 | -1.47 | -2.27 | 0.80 | 0.92 |
| CAAN2 | -1.95 | -2.15 | 0.21 | 0.99 |
| CACO | NA | NA | NA | NA |
| CAIN3 | -0.08 | 0 | -0.08 | 1 |
| CAIN4 | 0.42 | -1.47 | 1.89 | **0.0013** |
| STBR | 1.77 | 0.60 | 1.17 | 0.25 |
| STDI | 0.89 | -0.60 | 1.49 | **0.012** |
| STDR | 0 | -0.42 | 0.42 | 0.95 |
| STGL | 1.61 | -0.51 | 2.12 | **0.0003** |
| STIN | -0.34 | -0.99 | 0.65 | 0.75 |
| STPO | 1.61 | 1.33 | 0.27 | 0.99 |
| STTO | 0.51 | -1.21 | 1.72 | **0.0029** |

**Table S9.** Results from comparisons of germination rates between historical (Oct-2) and contemporary (Oct-30) onset of seasonal precipitation. Linear models were fit with germination rates as the response variable and the timing of rainfall onset (cohort) as the predictor variable. Comparisons of estimated marginal means between rainfall onset dates were made using the emmeans package (Lenth 2022). Least-squares means are reported for Oct-2 and Oct-30, their Contrast value, and Tukey adjusted p-values of the contrasts. P-values in bold are significant. No comparison was made for CACO due to lack of germination in the Oct-30 rainfall event.

| **Species** | **Oct-2** | **Oct-30** | **Contrast** | **P-value** |
| --- | --- | --- | --- | --- |
| CAAM | 0.11 | 0.08 | 0.02 | 0.92 |
| CAAN1 | 0.18 | 0.14 | 0.04 | 0.85 |
| CAAN2 | 0.20 | 0.13 | 0.07 | 0.37 |
| CACO | NA | NA | NA | NA |
| CAIN3 | 0.24 | 0.17 | 0.07 | **< 0.0001** |
| CAIN4 | 0.18 | 0.17 | 0.01 | 0.99 |
| STBR | 0.13 | 0.11 | 0.02 | 0.22 |
| STDI | 0.31 | 0.18 | 0.13 | **< 0.0001** |
| STDR | 0.12 | 0.11 | 0.01 | 0.99 |
| STGL | 0.14 | 0.11 | 0.03 | 0.19 |
| STIN | 0.21 | 0.13 | 0.08 | **< 0.0001** |
| STPO | 0.20 | 0.14 | 0.06 | **< 0.0001** |
| STTO | 0.30 | 0.16 | 0.13 | **< 0.0001** |

**Table S10.** Results of *post hoc* analyses from a Pearson’s Chi-squared test with three contingencies of germination timing, Year 1, Year 2, and Never. Bonferroni adjusted p-values in bold are significant.

| **Species** | **Value** | **Year.1** | **Year.2** | **Never** |
| --- | --- | --- | --- | --- |
| CAAM | Residuals | -10.67 | -3.75 | 12.55 |
| CAAM | p values | **0** | **0.0069** | **0** |
| CAAN1 | Residuals | -10.09 | 13.04 | 2.87 |
| CAAN1 | p values | **0** | **0** | 0.16 |
| CAAN2 | Residuals | -10.79 | 9.06 | 5.72 |
| CAAN2 | p values | **0** | **0** | **0** |
| CACO | Residuals | -1.31 | -1.51 | 2.11 |
| CACO | p values | 1 | 1 | 1 |
| CAIN3 | Residuals | 2.85 | -2.07 | -1.68 |
| CAIN3 | p values | 0.17 | 1 | 1 |
| CAIN4 | Residuals | -2.35 | -4.79 | 4.92 |
| CAIN4 | p values | 0.73 | **< 0.0001** | **< 0.0001** |
| STBR | Residuals | 9.32 | -3.33 | -7.38 |
| STBR | p values | **0** | **0.03** | **0** |
| STDI | Residuals | 0.99 | -2.28 | 0.25 |
| STDI | p values | 1 | 0.88 | 1 |
| STDR | Residuals | 2.85 | -4.59 | -0.32 |
| STDR | p values | 0.17 | **< 0.0001** | 1 |
| STGL | Residuals | 0.42 | 10.32 | -6.01 |
| STGL | p values | 1 | **0** | **0** |
| STIN | Residuals | -3.62 | -2.91 | 5.15 |
| STIN | p values | **0.01** | 0.14 | **< 0.0001** |
| STPO | Residuals | 11.51 | -2.91 | -9.77 |
| STPO | p values | **0** | 0.14 | **0** |
| STTO | Residuals | 10.24 | -5.01 | -7.38 |
| STTO | p values | **0** | **< 0.0001** | **0** |

**Table S11.** Principal component analysis of average germination season (September - December) climate for 1991-2015 for species’ location. First two principal component axes (PCs) illustrated in Figure 1B. Climate variables include precipitation (PPT), minimum and maximum temperature (Tmin, Tmax, respectively), as well as variability in precipitation (coefficient of variation, PPT_CV) and temperature (standard deviation of Tmin and Tmax, Tmin_SD and Tmax_SD respectively). Data source was Flint and Flint, (2014).

| **Variable** | **PC1** | **PC2** |
| --- | --- | --- |
| PPT | -0.49 | -0.31 |
| PPT_CV | -0.07 | 0.76 |
| Tmin | 0.44 | 0.25 |
| Tmin_SD | 0.29 | 0.31 |
| Tmax | -0.47 | -0.31 |
| Tmax_SD | 0.51 | -0.28 |
| Cumulative Proportion | 0.44 | 0.67 |

**Figure S1**. Location of populations used in this study within the species’ range-wide temperature and precipitation space based on herbarium records from the Consortium of California Herbaria. A principal component analysis including all species was performed of average germination season (September - December) climate for 1991-2015. Principal component scores were subset for each species and the first two principal components (PCs) are illustrated on the x and y axes, respectively. Climate variables include precipitation, minimum and maximum temperature, as well as variability in precipitation (coefficient of variation) and temperature (standard deviation). Climate data was sourced from the California Basic Characterization Model (Flint and Flint 2014). Population(s) included in this study are shown as a black triangle. For STTO, points are colored by elevation since this species spans a wide elevational range, but our study population is from low elevation (511 m).

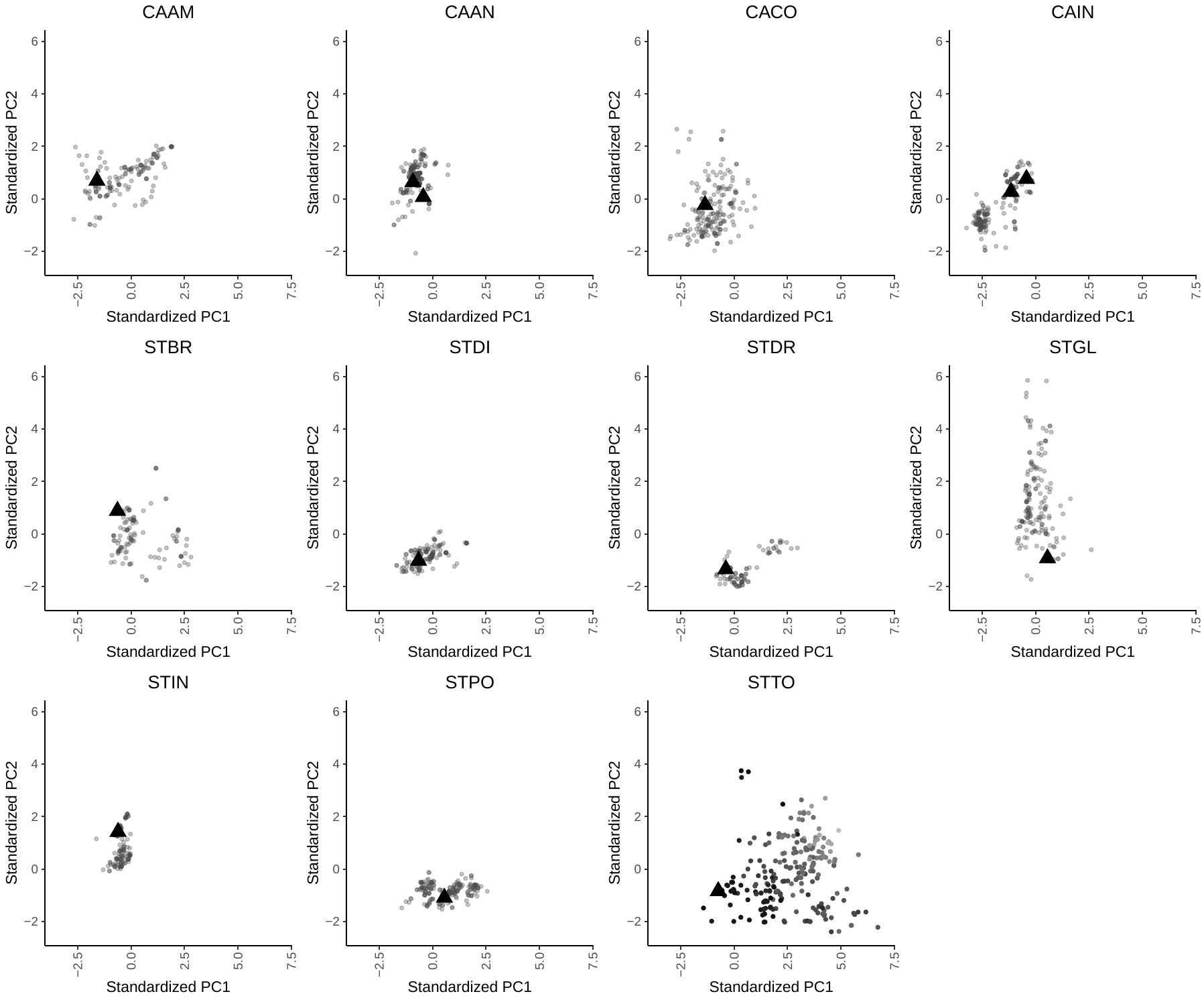

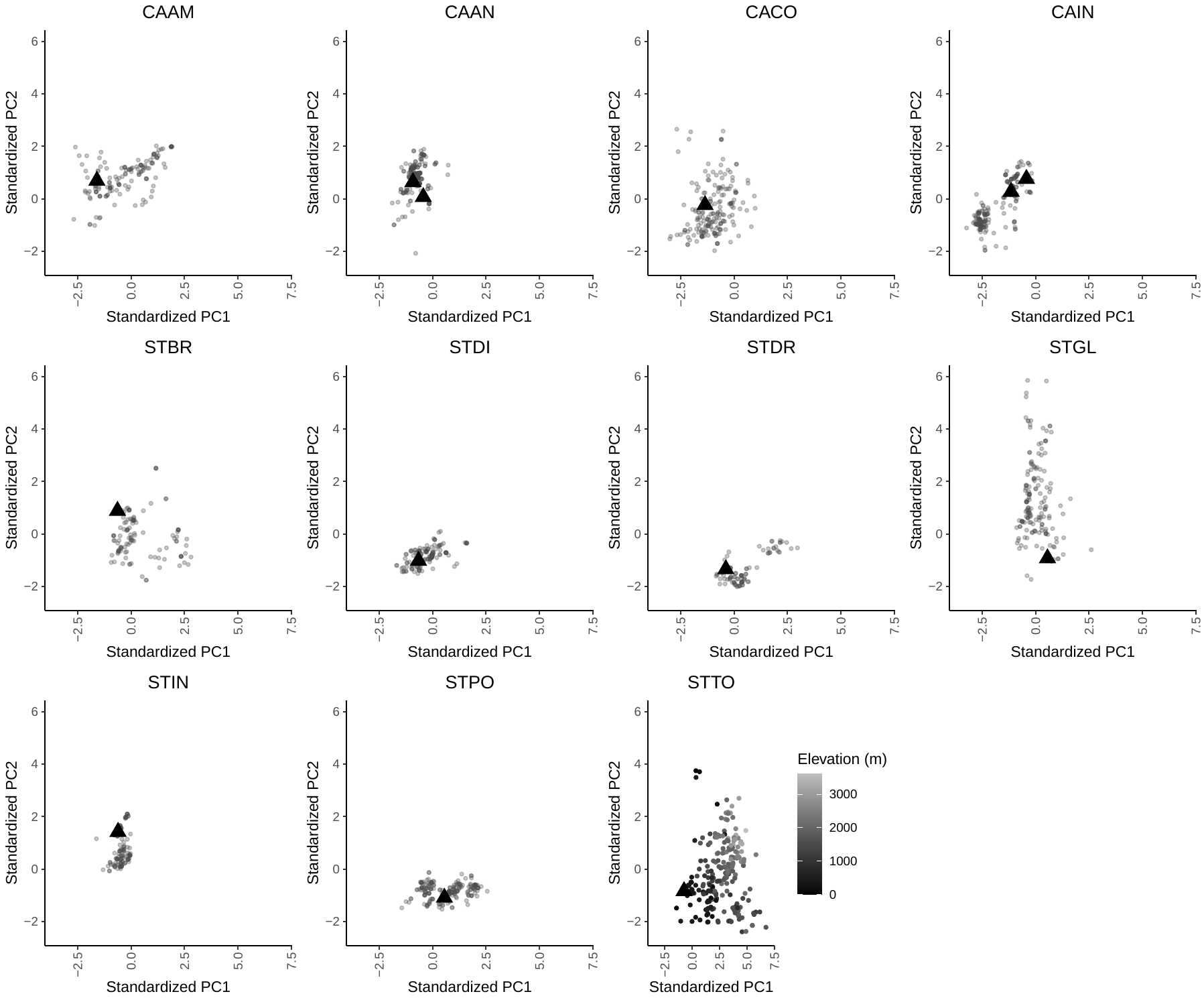
